# Identification of compounds producing non-visual photosensation via TRPA1 in zebrafish

**DOI:** 10.1101/2020.06.10.111203

**Authors:** Darya Cheng, Matthew N McCarroll, Jack C Taylor, Taia Wu, David Kokel

## Abstract

TRPA1 receptors sense chemical irritants, but they do not normally respond to light. Previous studies have identified compounds that confer photosensitivity onto vertebrate TRPA1. However, the pharmacology of TRPA1-mediated non-visual photosensation remains poorly understood. To identify novel compounds that affect this process, we screened a large chemical library for compounds that increased light-elicited motor activity in larval zebrafish. We found structurally diverse hit compounds that were photoreactive and produced specific behavioral phenotypes. A subset of these compounds required functional TRPA1 to produce behavioral phenotypes in vivo. These findings provide novel prototype compounds for controlling TRPA1 with light and improve our understanding of non-visual TRPA1-mediated photosensation.

## INTRODUCTION

TRPA1 receptors are members of the transient receptor potential (TRP) channel family^1^. They are tetrameric ion channels^2^ with cytoplasmic ankyrin repeats that mediate sensitivity to thermal and chemical stimuli^3^. TRPA1 channels respond to a wide variety of reactive compounds including mustard oil^4^, industrial electrophiles^5^, drug metabolites^6^, and other compounds^1,7^. They mediate the inflammatory actions of environmental irritants and proalgesic agents^8^. In addition, they mediate cough^9^, inflammation^5^, and airway irritation^10^. A gain-of-function mutation in TRPA1 causes familial episodic pain syndrome^11^. These functions are important for diverse animal behaviors.

A relatively small subset of TRPA1 isoforms also respond to light. For example, in specific species of snakes, TRPA1 channels sense infrared light, aiding in prey capture^12^. In human melanocytes, TRPA1 channels respond to UV light, increasing protective pigmentation^13^. Furthermore, TRPA1 homologs are required for extraocular-light avoidance in planaria^14^. Additionally, TRPA1 channels mediate light-induced feeding deterrence^15^ and UV light avoidance^16^ in drosophila. However, despite these examples, most TRPA1 isoforms, including human and zebrafish isoforms, do not normally respond to light^3^.

Understanding TRPA1-mediated photosensation is important for understanding non-visual photo-behaviors. In humans, non-visual photo-behaviors include circadian rhythms, alertness^17^, and the pupil reflex^18^. In other species, more diverse non-visual behaviors are observed, including phototaxis in zebrafish^19^, migration in birds^20^, and startle reflexes in insects ^21^. In humans, all visual and non-visual behaviors are mediated by opsins^22^. In non-human species, non-visual behaviors are mediated by at least four additional types of photoreceptors including cryptochromes^23^, photoactivated adenylyl cyclases (PACs)^24^, gustatory related receptors (GRRs)^25^ and TRPA1. TRPA1-mediated non-visual photosensation may represent a potential ancestral function of this gene^14^.

Previous studies have identified an exogenous small molecule that enables non-visual TRPA1-mediated photosensation in vertebrates. This small molecule, optovin, is photoactivated by relatively short wavelengths of visible light^26^. Its activity is both light-dependent and rapidly reversible. It acts on various TRPA1 isoforms including human, mouse, and zebrafish^26^. As a result, it enables light-mediated control of transgenic cardiac cells^27^ and of endogenous TRPA1-expressing neurons. The photoactive properties of optovin have been linked to its rhodanine ring, a chemical moiety known to absorb light^28^. Several structural analogues of optovin have been identified that also produce similar phenotypes^27^. These data suggest that there may exist other compounds that are functionally related to optovin that also produce non-visual photosensation via TRPA1.

Despite this progress, few compounds beyond optovin have been identified that activate TRPA1 *in vivo* via photoreactivity. To identify such compounds, we took a behavior-based drug profiling approach in zebrafish. Specifically, we screened a large chemical library for compounds that increased motor activity in response to light stimuli. This screen identified a set of structurally diverse hit compounds that were both photoreactive and produced specific behavioral profiles. A subset of these behavioral phenotypes required functional TRPA1. Together, these data improve our understanding of non-visual photosensation. In addition, they provide structurally diverse prototype compounds that may enable researchers to control endogenous TRPA1 with light.

## RESULTS

### Optovin produces a distinct behavioral profile

Previous studies of how optovin affects behavior have primarily focused on early stage zebrafish embryos^26^. In embryos, the optovin response includes two spikes of motor activity-a large initial spike followed by a smaller subsequent spike of activity^26^. These motor activity spikes are thought to correspond to C-bend behaviors and swimming behaviors, respectively. However, less is known about optovin activity on zebrafish larvae, which exhibit more complex behaviors than embryos, and are a powerful system for studying the vertebrate CNS. Therefore, it is important to characterize optovin activity in zebrafish larvae.

To determine how optovin affects zebrafish larvae, we treated larvae with various TRPA1 ligands, tested them with different stimulus batteries, and analyzed their resulting behavioral profiles. First, larvae were treated with optovin for one hour, and, following a three-minute delay, exposed to a single pulse of light (10 min). Prior to light exposure, the optovin-treated animals demonstrated a significantly lower baseline activity level than DMSO-treated animals. Following activation of the stimulus, the response of optovin-treated animals could be divided into three phases. In the first phase, P1, onset of the stimulus elicited a sharp spike of motor activity, represented as the motion index, or MI (**Fig. 1a, b**). The MI during this spike increased approximately 8-fold over baseline, and lasted for approximately 6s. (**Fig. 1a, b, Table S1**). In the second phase, P2, motor activity spiked again, increasing approximately 3-fold over baseline and lasting for approximately 1 minute (**Fig. 1a, b, Table S1**). In the third phase, P3, motor activity remained slightly higher than the optovin baseline for the duration of the stimulus (**Fig. 1a, b**). Following the stimulus, motor activity reverted to baseline (**Fig. 1a, b**). The P1 and P2 motor activity spikes were elicited only by blue (460nm) and violet (395nm) wavelengths, but not by longer wavelengths including green (523nm) and red (660nm) (**Fig. 1c, Fig. S1**). Unlike in optovin-treated animals, the P1 and P2 spikes were not observed in control larvae treated with the solvent alone. In P1, the control response only increased 2-fold — substantially less than the 8-fold increase observed in optovin-treated animals (**Fig. 1a, b**). In P2, the control response did not include a broad activity spike, unlike the optovin response. In P3, the control response did not decline (**Fig. 1a, b**). Another difference was that the control response included a distinct lights-off response at the end of P3 that was not observed in optovin-treated animals (**Fig. 1a)**.

**Figure 1.**
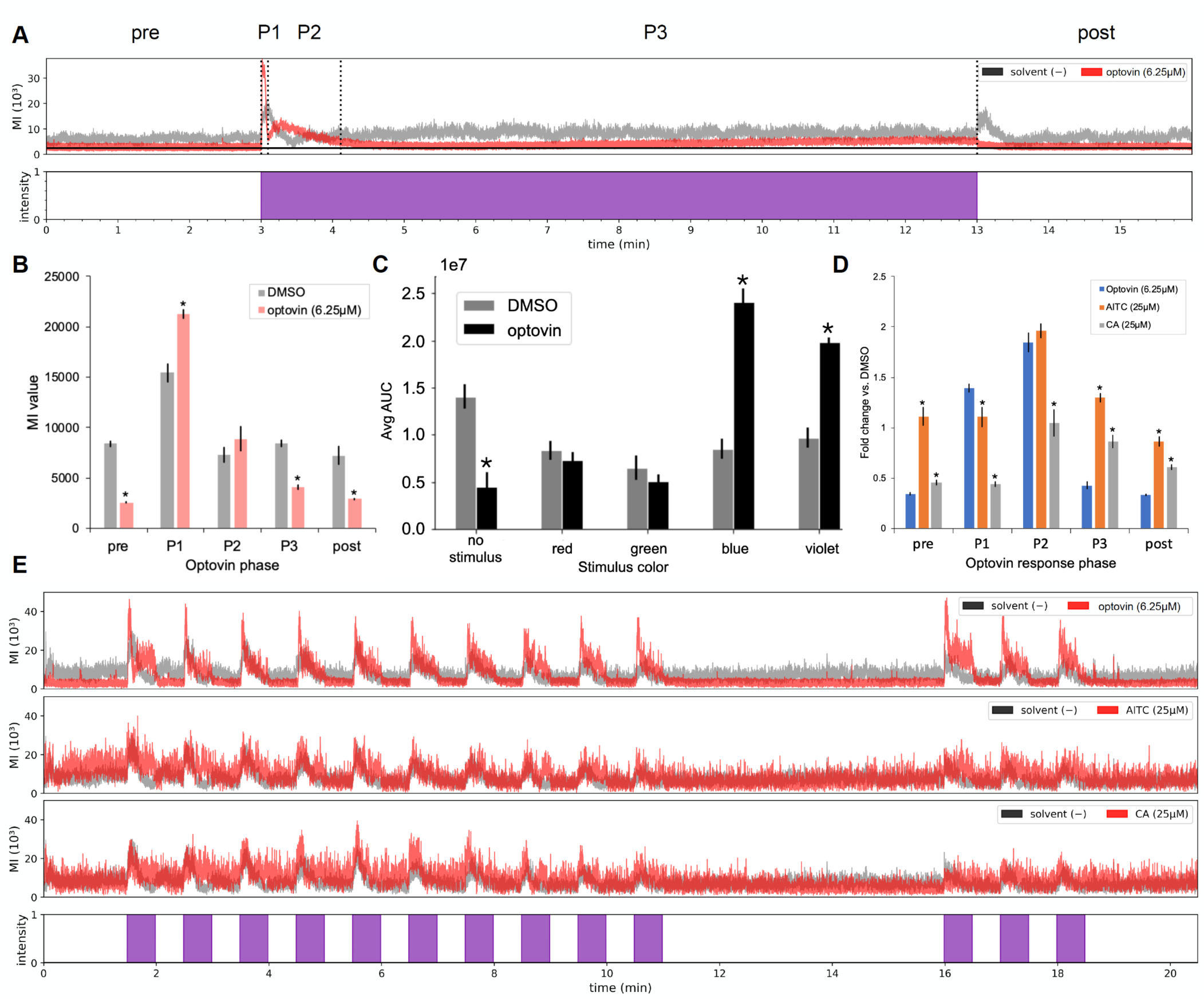
Optovin increases light-dependent motor activity spikes in zebrafish larvae. (A) Optovin treated zebrafish exposed to violet light (395nm) display three phases of activity that are distinct from control-treated larvae, shown here in a motor activity plot. Activity is shown as the motion index (MI) on the y-axis, with time on the x-axis. Colored bars below the motor activity plot indicate stimuli turning on (colored bar) and off (white bar); colors correspond to stimulus wavelength. (B) Bar plot showing average MI values (y-axis) for each optovin phase (x-axis). Data shown as mean ± SEM of n = 72 optovin-treated wells (red) and 40 DMSO-treated wells (gray) (8 fish/well); * p<0.05. (C) Bar plot showing AUC (y-axis) of P1 and P2 spikes in optovin-(black) and DMSO-(gray) treated larvae following a 1 minute light stimulus. In optovin treated fish, P1 and P2 motor activity spikes following were elicited only by blue (460nm) and violet (395nm) wavelengths. Red (660nm), green (523nm), and no stimulus conditions did not elicit motor activity. Data shown as mean ± SEM of n= 6 wells per stimulus for both optovin and DMSO (8 fish/well); * p<0.05. (D) Bar plot showing activity fold change vs. DMSO (y-axis) of optovin-(blue), allyl isothiocyanate-(AITC, orange), and cinnamaldehyde-(CA, gray) treated larvae during each of the optovin-defined phases (x-axis). Data shown as mean ± SEM of n= 8 wells (8 fish/well); * p<0.05. (E) Representative activity traces for larvae treated with the indicated compounds (red) or control (grey). AITC, allyl isothiocyanate; CA, cinnamaldehyde. Data shown as mean of n= 3 wells (8 fish/well).

We observed several differences between the response to optovin and more canonical TRPA1 ligands including allyl isothiocyanate (AITC) and cinnamaldehyde (CA). One difference was that in both AITC-treated and CA-treated animals, motor activity throughout the assay, including during the P2 phase, broadly increased (**Fig. 1d)**. However, this broad increase in activity did not include the distinct P1 spikes observed in optovin-treated animals (**Fig. 1e)**. Another difference was that inter-stimulus activity levels increased in both AITC-treated and CA-treated animals (**Fig. 1d, e**). By contrast, in optovin-treated animals, inter-stimulus activity levels decreased (**Fig 1d, e)**. A third difference was that, in AITC and CA treated animals, multiple stimuli did not elicit multiple P1 or P2 spikes (**Fig. 1d, e)**. By contrast, in optovin-treated animals, multiple stimuli did multiple elicit P1 spikes. Together, these data suggest these P1 and P2 motor activity spikes may represent features of a specific behavioral profile that correlates with Trpa-mediated non-visual photosensation. The reason is that these P1 activity spikes specifically occurred in optovin-treated zebrafish larvae but were not observed in untreated larvae, or in larvae treated with other TRPA1 ligands, such as AITC and CA. Alternatively, this behavioral profile may be a random coincidence with little predictive power for identifying novel TRPA1 ligands. To distinguish between these alternatives, we analyzed a larger number of compounds.

### Large-scale chemical screening

In previous studies, large-scale behavior based chemical screens in zebrafish have successfully identified diverse classes of neuroactive compounds related to anesthesia^29^, sleep^30^, fear^31^, feeding^32^, and antipsychotics^33^. Despite this progress, it remains unclear if novel compounds that produce TRPA1-mediated non-visual photo sensation can be identified based on their behavioral profiles. To answer this question, we analyzed behavioral data collected from a large-scale chemical screen (**Fig. S2**).

In this screen, the chemical library included 30,656 structurally diverse compounds. These compounds were profiled on zebrafish larvae using a battery of behavioral phenotyping assays that included a variety of stimuli, including light-based stimuli. The primary screening hits, defined as any compound that substantially increased motor activity in response to light, included 269 potential hit compounds including 54 false positive solvent-treated controls (**Fig. 2a**). These potential hits were manually reviewed to prioritize the most promising hit compounds and remove false positives due to technical errors, yielding 99 primary hits (**Fig. S2**). From these hits, 6 optovin analogues were removed, and a subset of 93 primary hits were reordered and retested (**Fig. S2**). Of these 93 primary hit compounds, 34 (36.56%) reproducibly increased light-induced motor activity in vivo, and 17 compounds were selected as final hits following dose-response testing. An additional four compounds were included based on manual analysis identifying them as potential hits of interest for a final total of 21 hit compounds (**Fig. 2b, Fig. S2**).

**Figure 2.**
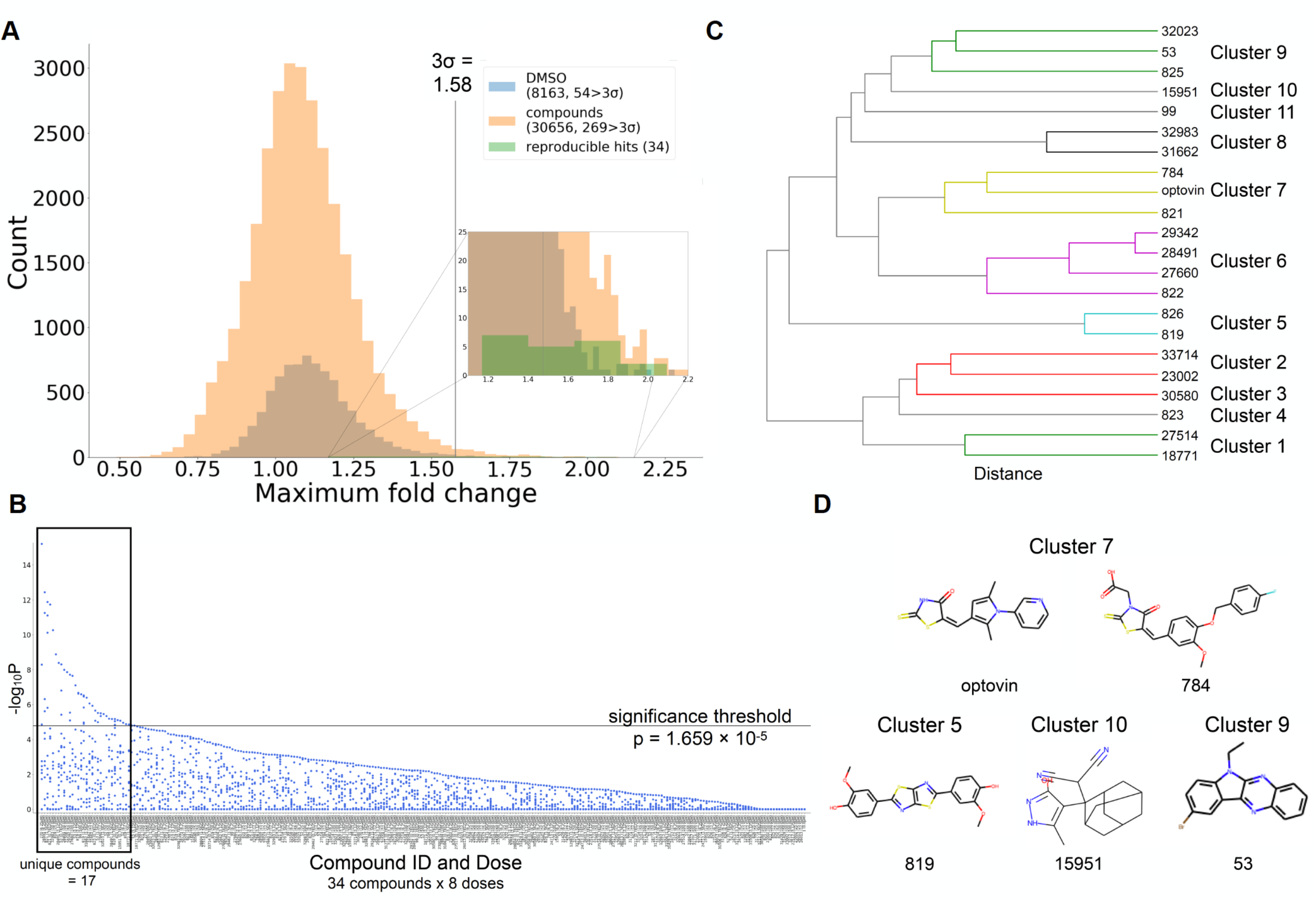
Large-scale screening identifies optovin-related hit compounds. (A) Two chemical libraries, containing a combined 30,656 structurally diverse compounds and 8,163 negative DMSO controls, were tested against a battery containing a variety of light-based stimuli. Fold change in activity versus DMSO was calculated for each compound for stimuli of interest. Compounds that had at least one measured value with a fold change greater than 3σ (1.58) were selected for further retesting, from which 34 were selected for dose-response testing. Data shown as n = 1 well per compound and 8163 wells DMSO; 8 fish/well. (B) Scatter plot showing -log_10_P values for selected stimuli of interest for each compound-treated well compared against the corresponding stimulus in DMSO-treated wells. The 34 compounds selected for dose-response analysis were tested at 2-fold intervals from 0.78125μM to 100μM against the screening battery. Seventeen unique compounds exceeded the significance threshold (horizontal line). n = 6 wells/test compound/dose, 368 wells DMSO, 118 wells optovin, and 96 wells with no treatment; 8 fish/well. Significance threshold corrected using Bonferroni correction. *p < 1.659×10^−5^ (C) Dendrogram depicting hierarchical clustering of hit compound structures. (D) Examples of hit compound structures.

Structural analyses revealed that the validated hit compounds included a wide variety of structures. These compounds clustered into at least 11 structural classes (**Fig. 2c, Fig. S3, Table S2**). Unlike optovin, most of the novel hit compounds did not contain a rhodanine ring except compound 784. However, there were several other features of note. Though they did not cluster together, compounds 15951 (cluster 10) and 23002 (cluster 2) both have an adamantane group. Additionally, compounds 819 and 825 in cluster 5 both have a central, fused double thiazole along with distinct symmetry in each compound. Likewise, compounds in cluster 6 all share a thienopyridine. Together, these data suggest that, like optovin, a subset of these novel compounds may also be photoactivated TRPA1 ligands, as the validated hit compounds reproducibly produce light-elicited behavioral excitation. Interestingly, the hit compounds included a variety of chemical structures, raising questions about their potential mechanisms of action and photoreactivity.

### Photoactivity of hit compounds greatly varies

Previous studies have revealed that aspects of TRPA1-mediated photo-sensation involve photoreactivity. For example, TRPA1 is activated by photosensitizing agents *in vitro*^*34*^. Furthermore, optovin is a photoreactive compound^26^. Thus, we hypothesized that a subset of hit compounds would also be photoreactive. To answer this question, we measured a variety of photochemical properties including their absorbance, photostability after multiple irradiations, fluorescence, and phototoxicity.

The absorbance studies revealed that most compounds displayed peak absorbance between wavelengths ∼310 and 410nm (**Fig. 3a, Fig. S4**). For example, 99, 784, and 28491 displayed peak absorbance at 390nm, 410nm, and 350nm, respectively **(Fig. 3a, Fig. S4)**. Several compounds produced two absorbance peaks including 53, 825, and 27660 **(Fig. 3a, Fig. S4)**. By contrast, none of the novel hit compounds had a maximum absorbance above 410nm (**Fig. 3a, Fig. S4**), while six compounds (15951, 18771, 27514, 30580, 31662, 33714) did not display absorbance at any wavelength tested. Optovin absorbance, by comparison, occurs at 470nm, which is an outlier compared to the absorption range of the hit compounds (**Fig. 3a, Fig. S4**). The DMSO-treated negative controls displayed no absorption at any wavelength in the tested range (**Fig. 3a, Fig. S4**).

**Figure 3.**
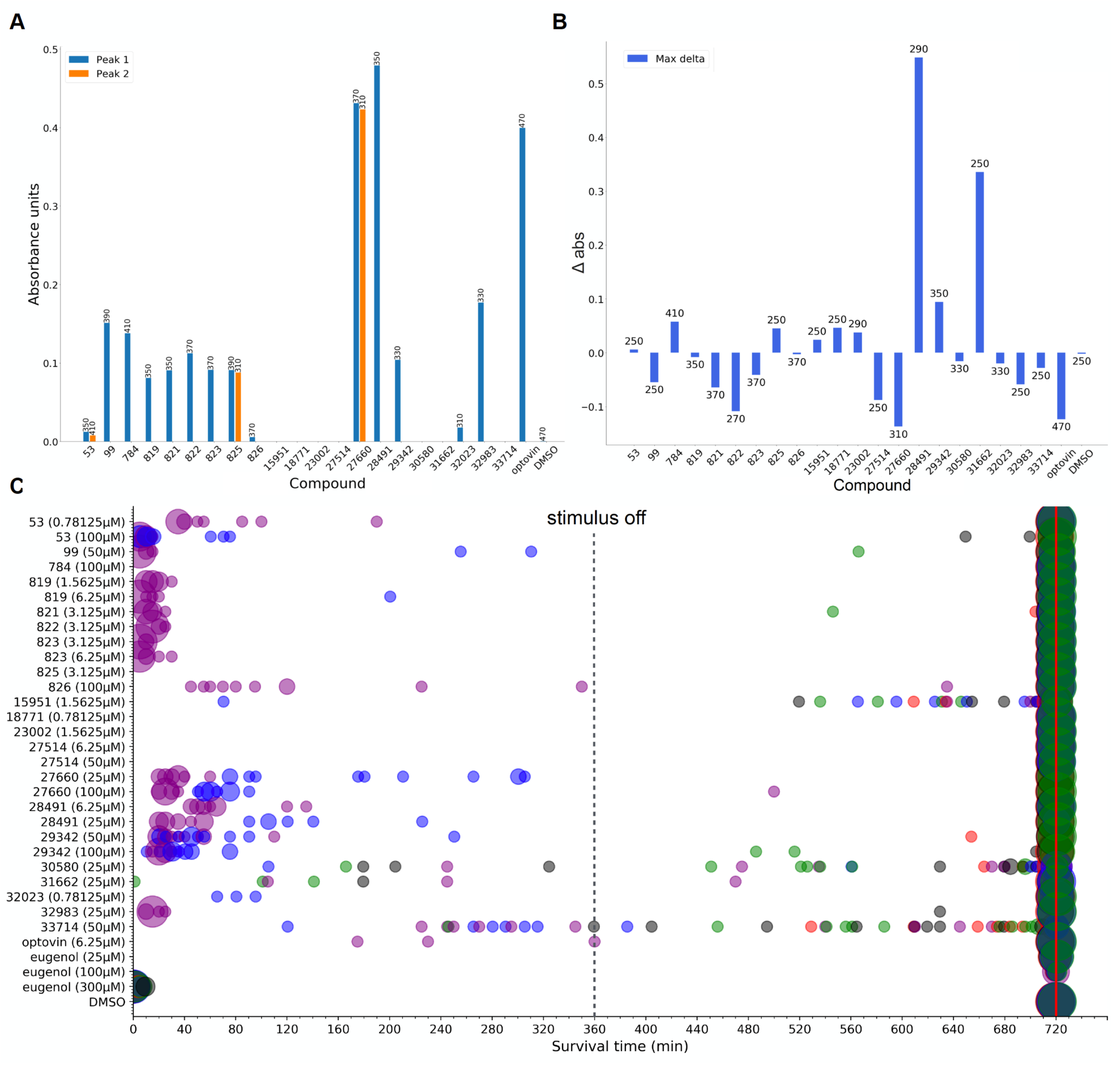
Hit compounds demonstrate photoreactivity. (A) Bar plot showing the magnitude (y-axis) of light absorbance of hit compounds. Where more than one absorbance peak was detected, the bar for the secondary peak is colored in orange. Absorbance wavelengths (nm) are labeled at the top of each bar. (B) Bar plot showing the maximum change in absorbance (y-axis) following multiple irradiation. Following an initial baseline absorbance measurement, compounds were exposed to violet light for varying lengths of time between 5 and 1200s (2 min.), after which absorbance was measured again. Max delta refers to the greatest change in absorbance at one wavelength between any two timepoints. Absorbance wavelengths (nm) are labeled at the top of each bar. (C) Plot depicting the survival time of hit compound-treated larvae exposed to varying wavelengths of light. The activity of treated fish was tracked for 12 hours. During the first six, larvae were exposed to light stimulus, after which activity was tracked for an additional six hours. Marker color indicates stimulus wavelength (no stimulus condition indicated by black). The size of the marker indicates the number of wells, with the position of the marker along the x-axis indicating the time at which those wells were considered to have died. Time of death was determined by identifying the last 5 minute window in which the standard deviation of activity was equal to or less than the overall standard deviation of activity in the 300μM eugenol-treated positive controls. Wells that did not meet this criterion within the 12 hour period were considered to have survived. Data shown as mean of n = 12 wells per stimulus per compound (8 fish/well).

As a measure of photostability, we determined if the compounds changed absorbance after multiple irradiations. In 17 of 21 hit compounds, the maximum change in absorbance was equal to or less than 0.10 absorbance units, a change that was relatively minor (**Fig. 3b, Fig. S5, Table S3**). For example, 7 compounds (including 53, 784, 825, 15951, 18771, 23002, 29342) increased absorbance by 0.10 absorbance units or less (**Fig. 3b, Fig. S5, Table S3**). Similarly, 10 compounds (including 99, 819, 821, 823, 826, 27514, 30580, 32032, 32983, 33714) decreased absorbance by less than 0.10 absorbance units (**Fig. 3b, Fig. S5, Table S3**). By contrast, multiple irradiations greatly increased the absorbance of compounds 28491 and 31662, rising by 0.55 and 0.34 absorbance units, respectively, while absorbance in compounds 822 and 27660 was reduced by 0.11 and 0.14, respectively (**Fig. 3b, Fig. S5, Table S3**). Comparatively, the 0.12 absorbance unit reduction of peak absorbance in optovin occurred at 470nm, while DMSO demonstrated no change (**Fig. 3b, Fig. S5, Table S3**).

Fluorescence studies revealed that most hit compounds were not strongly fluorescent (**Table 1**). For example, 14 compounds (53, 784, 819, 825, 826, 15951, 18771, 23002, 27514, 29342, 30580, 32023, 32983, 37714) had excitation/emission wavelengths of 350nm/395nm (**Table 1**). By contrast, a subset of compounds showed measurable emissions. For example, compounds 821 and 822 emitted in the blue range (455nm and 475nm, respectively) (**Table 1**). Compounds 99, 823, 27660 (all 535nm) and 28491 (515nm) emitted in the green range (**Table 1**). One outlier was compound 99, which displayed a maximum excitation wavelength at 410 nm (**Table 1**). Importantly, neither optovin nor DMSO were strongly fluorescent, displaying maximum excitation/emission at 350nm/395nm (**Table 1**).

**Table 1.**
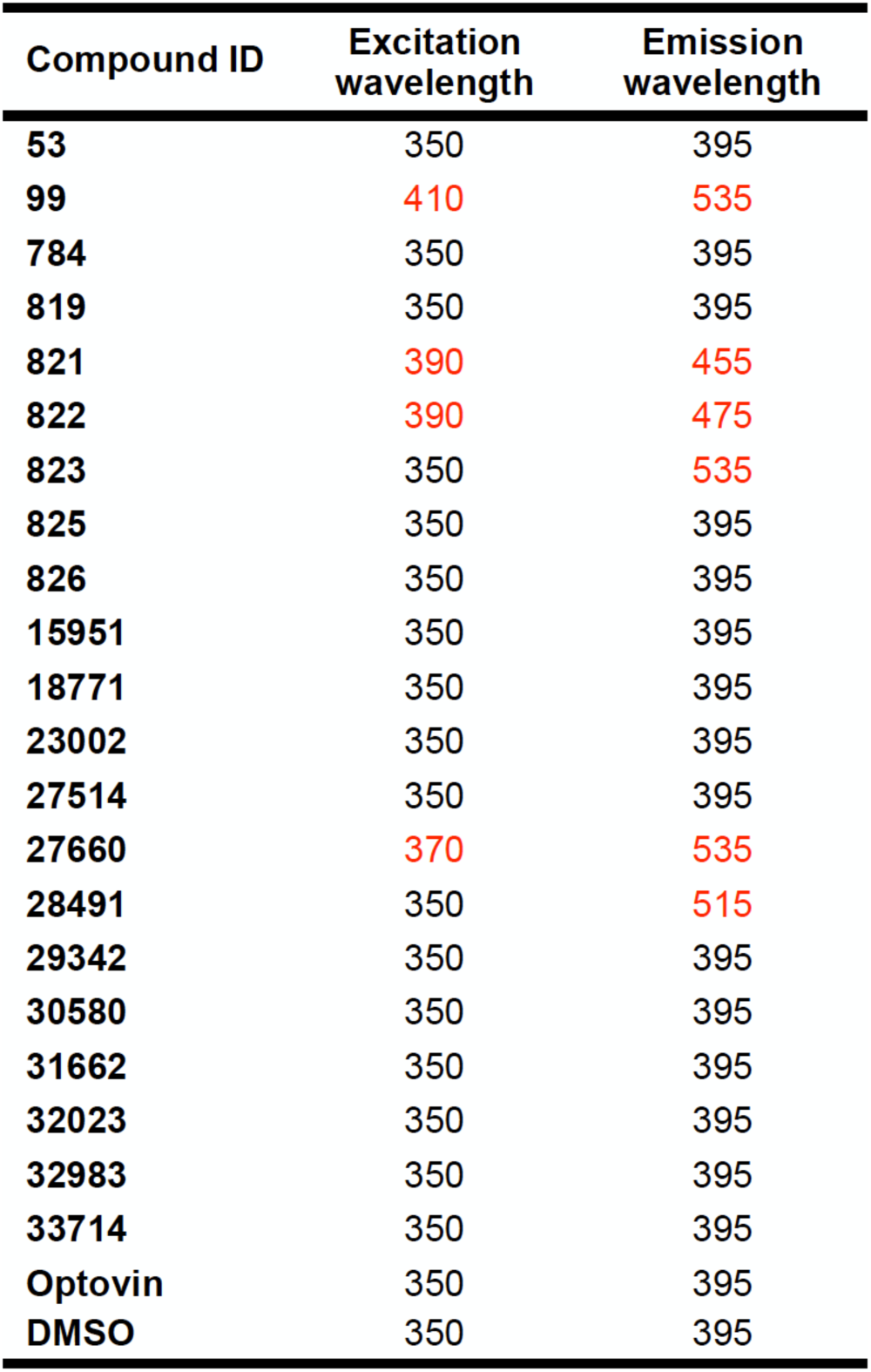
Excitation/emissions experiments showed that most compounds are not strongly fluorescent. For each compound, the maximum excitation and emission wavelengths were identified. Outliers that excite/emit at wavelengths other than 350nm/395nm are highlighted in red.

| Compound ID | Excitation wavelength | Emission wavelength |
| --- | --- | --- |
| 53 | 350 | 395 |
| 99 | 410 | 535 |
| 784 | 350 | 395 |
| 819 | 350 | 395 |
| 821 | 390 | 455 |
| 822 | 390 | 475 |
| 823 | 350 | 535 |
| 825 | 350 | 395 |
| 826 | 350 | 395 |
| 15951 | 350 | 395 |
| 18771 | 350 | 395 |
| 23002 | 350 | 395 |
| 27514 | 350 | 395 |
| 27660 | 370 | 535 |
| 28491 | 350 | 515 |
| 29342 | 350 | 395 |
| 30580 | 350 | 395 |
| 31662 | 350 | 395 |
| 32023 | 350 | 395 |
| 32983 | 350 | 395 |
| 33714 | 350 | 395 |
| Optovin | 350 | 395 |
| DMSO | 350 | 395 |

As another measure of photoreactivity, we determined if the compounds produced phototoxicity. To this end, we tracked the activity of treated fish for 12 hours. During the first six, larvae were exposed to light stimulus, after which activity was tracked for an additional six hours. Length of survival/time of death was calculated quantitatively by analysis of the MI traces. These phototoxicity studies revealed that many of the hit compounds (12/21) demonstrated some level of phototoxicity under violet and blue wavelengths (**Fig. 3c, Table S4-5**). For example, four compounds (including 53, 27660, 28491, and 29342) were significantly phototoxic under both violet and blue wavelengths, and seven compounds (including 99, 819, 821, 822, 823, 826, 32983) were significantly phototoxic under only violet wavelengths. Other compounds demonstrated a trend towards phototoxicity, though not reaching significant levels. These include 99 and 819, which have some blue wavelength toxicity in addition to their violet wavelength toxicity, and 32023, which has only blue wavelength toxicity (**Fig. 3c, Table S4-5**). Additionally, two compounds (53 and 28491) showed a dose-dependent sensitivity to different wavelengths, with higher doses conferring phototoxicity under blue wavelengths, while lower doses conferred phototoxicity only in the presence of violet wavelengths (**Fig. 3c, Table S4-5**). Phototoxicity most often occurred within the first two hours of light exposure and death rarely occurred under the green, red, or no stimulus conditions (**Fig. 3c, Table S4-5**). While some fish did die under red and green light conditions, these tended to be single replicates, and death occurred relatively late in the experiment (>8 hours). Two compounds (30580 and 33714) were significantly toxic under all conditions, including when no stimulus was present, while another two,15951 and 31662, trended towards general toxicity. Interestingly, in compound 30580, which was shown to be generally toxic, the presence of red light significantly increased survival above the no stimulus condition (**Fig. 3c, Table S4-5**). A small number of hit compounds (5/21) were not phototoxic under any conditions including 784, 825, 18771, 23002, and 27514. Likewise, optovin and low dose (25 and 100uM) eugenol were not phototoxic, though optovin demonstrated a trend towards phototoxicity under violet wavelengths. Importantly, solvent-treated negative controls were not toxic under any conditions, independent of stimulus wavelength. Furthermore, eugenol-treated (300uM) positive controls were toxic under all conditions, independent of stimulus wavelength (**Fig. 3c, Table S4-5)**.

Together, these photoreactivity data suggest that although a subset of hit compounds were photoreactive, this photoreactivity was difficult to measure by absorbance, multiple irradiation, and fluorescence. However, most of the hit compounds were phototoxic to some degree. These data suggest that many of the hit compounds are photoreactive, raising questions about their target and phenotypic specificity.

### Hit compounds cluster into four behavioral phenotypic classes

Previous research has shown that compounds with similar behavioral profiles tend to act via similar molecular mechanisms, a key principle of phenotype based genetic screens^35^. Therefore, we wondered if the hit compounds produced distinct behavioral sub-classes. To answer this question, we analyzed their behavioral profiles using a multi-step process. First, optimal doses for each compound were selected by identifying the concentration of each compound that produced the maximum increase in motor activity (**Fig. 2b**). At this optimal concentration, all the hits were then tested in a behavioral battery designed to emphasize potential phenotypic differences. To visualize these differences, the behavioral profiles were analyzed by t-SNE (**Fig. 4a**) and pairwise clustering (**Fig. 4b**), then inspected by a manual analysis (**Fig. 4c**).

**Figure 4.**
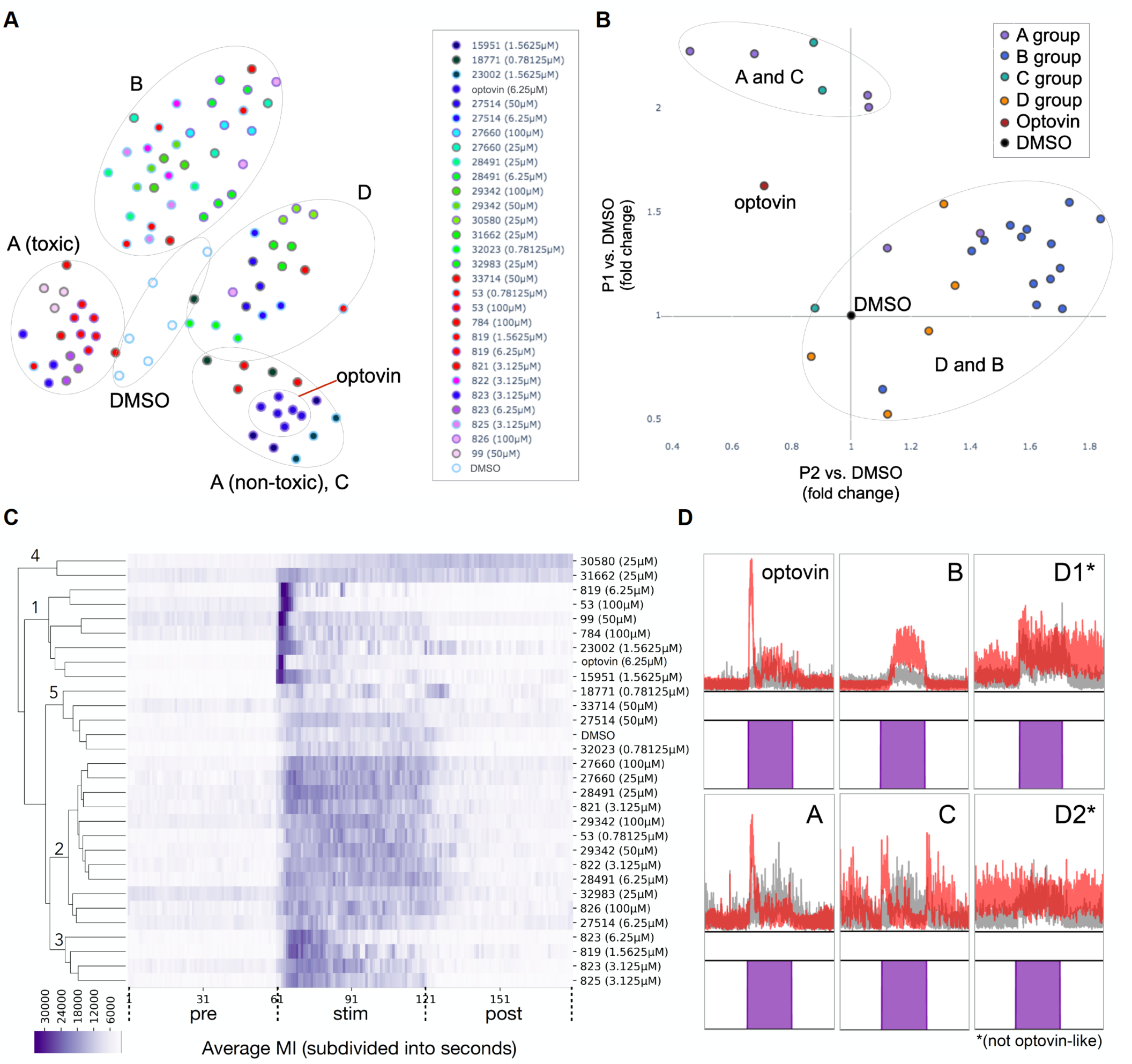
Hit compounds cluster into four phenotypic subgroups. (A) t-SNE analysis of full activity traces recorded from hit compound-treated larvae identifies clusters of compounds with similar phenotypes. Retroactive labelling following further analysis revealed that these groups closely correlated with the identified phenotypic clusters. Data shown as n = 6 wells optovin, 6 wells DMSO, 3 wells per hit compound; 8 fish/well. (B) Pairwise scatter plots identify the phenotypic features that best separate the groups identified in (A). Light responses for all compounds were first broken down into the optovin-defined phases (pre, P1, P2 and post, with P3 not represented in this test battery). Following pairwise comparisons, it was revealed that the P1 vs. DMSO and P2 vs. DMSO fold change measurements created the greatest separation between compounds. Retroactive labelling demonstrated that the presence of P1- or P2-like phases most strongly differentiated the A/C and B/D groups. Data shown as n = 6 wells optovin, 6 wells DMSO, 3 wells per hit compound; 8 fish/well. (C) Heatmap analysis and clustering further define phenotypic groupings. As labeled, branch 1, which includes optovin, is defined by a very strong initial spike of activity in the first few seconds of the stimulus, while branch 2 is characterized by increased activity throughout the stimulus. The phenotype observed in branch 3 has characteristics of both 1 and 2. Branches 4 and 5 display more anomalous phenotypes, with branch 4 displaying an increase of activity beginning during the stimulus and continuing through the post-stimulus period, while activity in branch 5 shows little consistency. Data shown as n = 6 wells optovin, 6 wells DMSO, 3 wells per hit compound; 8 fish/well. (D) Representative MI traces for the four phenotypic subgroups: A (most similar to optovin with a distinct P1 spike), B (purple light-sensitive with a P2-like response), C (on/off light responsive), and D (not optovin-like). Examples of each are shown here, including two examples for group D (red traces represent hit compounds, gray traces represent DMSO). Examples A, C, D1, D2: n = 3 wells; 5 fish/well. Examples for optovin, B: n = 3 wells; 8 fish/well.

These analyses revealed that the behavioral profiles could be classified into four main groups (Groups A-D), with DMSO-treated negative control wells clustered in a fifth group. The first group, Group A, included optovin and 4 additional compounds (53, 99, 784, 819) (**Fig. 4d, Fig. S7, Table S6**). These compounds produced phenotypes similar to the optovin response, characterized mainly by the P1 spike, which occurs at the onset of stimulus activity and is very distinctive (**Fig. 4c, d**) with the P2 spike sometimes but not always present (**Fig. 4b**). The P2 spike, if present, tends to be an elevated level of activity compared to background levels (during the no-stimulus portions of the battery), but is generally not greater than DMSO activity (**Fig. 4d, Fig. S7**). Referring back to the t-SNE analysis (**Fig. 4a**), category A compounds can be further split into “toxic” and “non-toxic” subgroups. However, because the B compounds likewise have varying levels of toxicity but are more closely grouped with each other rather than the toxic A compounds, this suggests that something beyond toxicity - most likely the P1 spike - separates the toxic A and B compounds.

The second group, Group B, included 13 compounds (**Fig. 4d, Fig. S8, Table S6**). These compounds produced phenotypes that were characterized by large P2 spikes, the absence of P1 spikes (**Fig. 4b, d**), and consistently elevated activity while the stimulus is ongoing. This response is usually 1.5- to 2-fold higher than DMSO and continues for the duration of the 1 minute light stimulus. Notably, in contrast to the category A response, the first few seconds of the response (the P1 “spike” seen in category A) are not increased compared to the rest of the response (**Fig. 4c, d**). Similar to category A, post-stimulus response is approximately the same or below DMSO-induced response.

The third group, Group C, included 3 compounds (**Table S6**) that displayed phenotypes very similar to those of optovin, but which can be distinguished by the presence of a second spike when the stimulus ends (**Fig. 4c, d, Fig. S9**). In this “on/off” (ON-OFF) response, when the stimulus is first activated, a sharp spike of activity occurs, similar to the Group A response. This similarity likely contributed to the inclusion of these compounds in the same clusters as category A compounds (**Fig. 4a-c**). However, unlike the Group A response, when the light stimulus is turned off, a second distinct activity spike is observed. Activity levels between the two spikes while the stimulus is ongoing are generally lower than DMSO, while post-stimulus activity is more variable than in both A and B, with occasional small spikes of activity. The three compounds that induce this behavior (15951, 18771, 23002) do so in response to all tested wavelengths of the light stimulus (**Fig. S6, Table S6**).

The fourth group, Group D, included 4 compounds that generally increased overall motor activity (**Fig. 4c, d, Fig. S10**). While the responses in Groups A, B, and C are to some extent optovin-like, Group D consists of non-optovin-like light-dependent behaviors. In general, while these behaviors are affected by the light stimulus, the immediate response to a stimulus is only slightly higher than in controls, but the slight increase in activity continues even after the stimulus ends. Additionally, inter-stimulus activity levels tend to be more variable than in the first three phenotypic categories.

Interestingly, of the compounds tested, some compounds (53, 819) induced different types of activity at different doses, where at low doses the response was more similar to the B group compound response, and at higher doses the response was more similar to those induced by the A group (**Table S6, Fig. S7, S8**). Furthermore, both long- and short-term toxicity of these compounds is mixed (**Fig. 3c**). In addition to violet light, blue, red, and green light stimuli were tested as well (**Fig. S6, Table S6**), revealing that similar patterns of behavior (A, B, C, or D) could be induced by these wavelengths, but not necessarily by the same compounds. For many compounds, blue light induced the same type of behavior in fish as violet light (**Fig. S6, Table S6**), but for others, blue light induced a different response or none at all (**Fig. S6, Table S6**). Red and green light in combination with hit compound treatments were less likely to elicit any response whatsoever, and those that did tended to induce category C or D behaviors (**Fig. S6, Table S6**). Since the hit compounds cluster into distinct phenotypic subclasses, these data suggest that each subclass may represent a different biological mechanism. For example, one of the most distinct clusters, the compounds that produce the Group C ON-OFF phenotype, may depend on different molecular targets than either optovin or the other hit compounds.

### A subset of hit compounds act via TRPA1b

Previous studies have shown that optovin produces non-visual TRPA1-mediated photosensation^26^. However, it remains unclear if any of the novel hit compounds depend on either vision or TRPA1. To answer this question, we analyzed the effects of mutations affecting TRPA1b as well as ATOH7, a gene that is required for the development of retinal ganglion cells and whose loss results in complete blindness^36^. Group A and B compounds, which induced activity most similar to that of optovin, were generally found to be dependent on TRPA1b (**Fig. 5a, b, c**). Conversely, group C activity is dependent on ATOH7, but not TRPA1b (**Fig. 5a, d, Fig. S11**). Group D activity did not depend on TRPA1b, but the dependence of this group as a whole on ATOH7 is unclear (**Fig. 5a, d, Fig. S11**). From the A group, P1 spiking was abolished in the TRPA1b mutants for four out of the five hit compounds. The last compound likewise followed a similar trend, though the change in response did not reach significance **(Fig. 5a, b)**. Similarly, the optovin response was abolished in TRPA1b mutants. Conversely, treatment using the A group compounds in ATOH7KO fish did not result in a reduced response (**Fig. S11**). Similarly, in Group B, P2 spiking was also abolished in TRPA1 mutants for six out of eleven compounds, but not in ATOH7 or WT controls **(Fig. 5a, c, Fig. S11)**. By contrast, for the two tested group C compounds, ONOFF was abolished in ATOH7 mutants. However, this did not occur in WT controls or in TRPA1 KOs, in which all three group C compounds were tested **(Fig. 5a, d, Fig. S11)**. Finally, in Group D, no compounds induced a significantly different response in TRPA1bKO fish compared to the WT fish. Two group D compounds were tested in the ATOH7 fish; of these, one showed a significant decrease in response **(Fig. 5a, d, Fig. S11)**. However, because only select compounds were tested in these fish, it is not possible to draw conclusions about the role of ATOH7 in each hit compound group, though certain trends can be observed. Together, these data suggest that compounds in Group A and B, like optovin, produce non-visual TRPA1-mediated photosensation. By contrast, the compounds in group C act on visual pathways.

**Figure 5.**
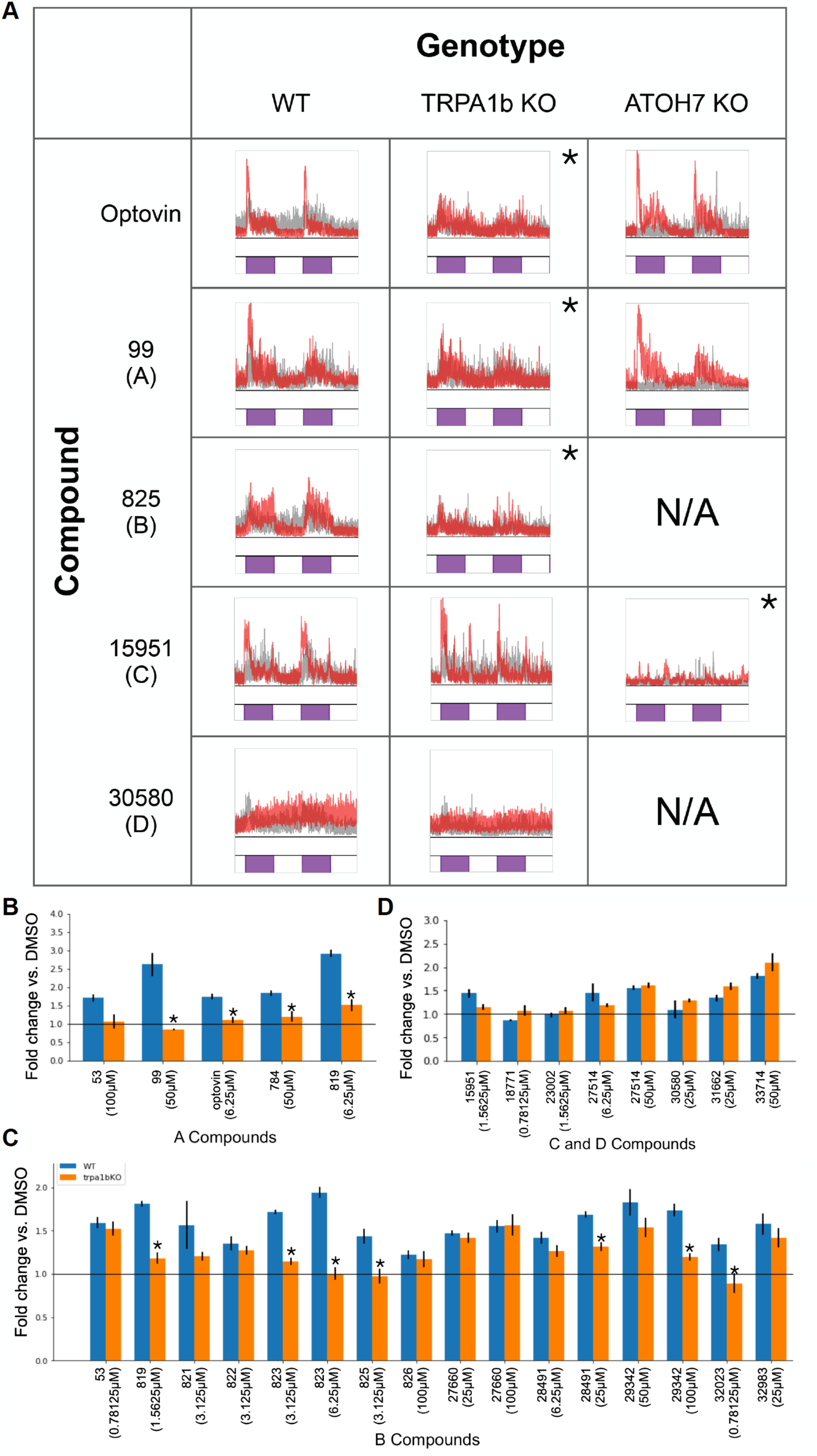
A subset of hit compounds act via the TRPA1b receptor. (A) Example traces illustrate the effect of hit compounds from different phenotypic groups in WT, TRPA1b KO, and ATOH7 KO fish compared to background-matched controls. Data shown as n = 3 wells; 5 fish/well. *p < 0.05. (B-D) Bar plots depict the response fold change of WT and TRPA1b KO fish treated with hit compounds against DMSO-treated background-matched larvae. (B) A group compounds; response measured during P1 phase. (C) B group compounds; response measured throughout the first stimulus (1 min.). (D) C/D group hit compounds; response measured throughout the first stimulus (1 min.). Data shown as n= 14 wells per compound per genetic background for DMSO and optovin, 3 wells per compound per genetic background for hit compounds, 2 wells per compound per genetic background for no treatment; 5 fish/well. Error bars represent SEM. *p<0.05.

## DISCUSSION

We demonstrate here that a wide variety of structurally diverse novel compounds can produce non-visual photosensation via TRPA1 in zebrafish. Evidence suggests that these compounds are photoreactive and depend on TRPA1b, and that they induce different patterns of behavioral phenotypes. These observations extend our understanding of the pharmacology of TRPA1-mediated non-visual photosensation beyond compounds structurally related to optovin

### Light responsive compounds demonstrate structural diversity

Although previous studies had identified optovin analogues^37^, it was unclear whether TRPA1-mediated non-visual photosensation also occurred with other, more structurally diverse compounds. Through this study, we have demonstrated that there are multiple novel compounds with structural and photochemical properties that differ greatly from those of optovin but are similarly able to induce motor activity in response to light. Previously, the photoactive properties of optovin have been linked to its rhodanine ring. However, the majority of photoactivatable compounds identified here do not contain this moiety, suggesting that a rhodanine ring may not be required to confer TRPA1-mediated non-visual photosensation.

### Hit compound phenotypes cluster into four distinct categories

Phenotypically, we observed that the compound responses to violet light tended to cluster into four main categories. Groups A and B comprised compounds that induce optovin-like phenotypes, with A defined by an initial sharp spike of activity (equivalent to P1 in optovin) and B defined by a more gradual increase in activity (equivalent to P2). This split was unexpected, as we initially believed that all optovin-like compounds would have a similar behavioral profile. One potential explanation for this is that the hit compounds - or, possibly, their light-activated configuration - may have varying affinities for TRPA1. Compounds with higher affinities would bind to and activate more channels, thus rapidly inducing a large amount of activity via the receptor, while compounds with lower affinities would not bind as frequently, resulting in fewer channel openings and therefore slower, more sustained activity that does not reach as high of a magnitude. Likewise, compound availability may be a limiting factor in receptor binding and may be affected by blood protein binding^38^ or distribution into the tissues^39^, or, if the compounds are acting on TRPA1 receptors in the brain, slow or limited passage across the blood-brain barrier^40^. This is supported by the observation that several of the compounds tested at multiple doses (53, 823) switch from B-type activity at the lower dose to A-type activity at the higher, where an increased dose would result in more of the compound being available to bind to receptors^41^. However, other compounds did not display A-type activity at any tested dose. In these cases, a possible explanation for B-type activity is that while the compounds may be readily available in their inactive state, they may undergo a slower conversion to the light-activated binding form, thus accounting for the relatively slower increase to maximum activity levels observed with B compounds. In addition to the A and B groups, two other groups were also identified.

Conversely, group C and D compounds did not induce optovin-like phenotypes. Group C compounds, despite their similarity to the A group due to the initial P1-like spike of the phenotype, also induced a secondary, smaller spike when the stimulus turned off, demonstrating a novel mode of light response. Group D consisted of compounds that induced phenotypes that were not similar to optovin and did not fit with the other categories. However, as these groups were not shown to act via TRPA1, it is difficult to speculate on their mechanisms.

### Optovin-like compounds demonstrate the greatest response to blue and violet wavelengths

In order to determine whether the light-induced phenotypes were specific only to violet light, we also tested the response to blue, red, and green wavelengths and categorized them using the same groups as for the violet-light response. In accordance with the observation that optovin also demonstrates a strong response to blue light, many of the A and B group compounds likewise respond to blue light stimuli, and for each compound the blue light activity type resembles that of the violet light. Considering all light dependent activity, we observed fewer responses to green and red wavelengths. This is in line with previous research^42^, as light in these wavelengths carry less energy than light in blue or violet, and therefore would possess less energy to contribute to structural changes. Because of this, it would be useful to perform a screen based on blue light activity to further probe the relationship between compound, wavelength, and receptor, and to identify whether blue light sensitivity exists independent of violet light sensitivity.

Interestingly, we observed that C and D compounds demonstrated activity in response to all four wavelengths. This suggests that the responses of both Group C compounds, whose activity is clearly light-dependent, and most Group D compounds, whose activity is more generally unspecific, are more of a generalized, non-wavelength-specific light response.

### Absorbance and fluorescence vary greatly

In addition to their structural and phenotypic features, we also investigated several photochemical properties of these compounds to determine if there were features that were common to all light-activated compounds. Though the wavelength at which absorbance occurred varied, optovin and all A and B group compounds exhibited some degree of absorbance. C and D compounds, however, did not demonstrate any absorbance. Together, this suggests that absorbance is required for light-dependent activation of TRPA1.

Prior exposure to violet light (for time periods ranging from 5 to 1200 seconds) had either no effect or reduced absorption on most hit compounds and optovin, likely due to the possibility of bleaching or non-reversible light-dependent structural changes. Interestingly, in some compounds (28491, 29342, 31662), prior exposure in fact greatly increased absorption, suggesting that the addition of light energy induced structural changes that increased the likelihood of energy absorption. The degree by which absorption changed did not seem to correlate to baseline absorbance (t0) (**Fig. 3b, Fig. S5**). Some compounds that demonstrated baseline absorbance showed very little change, while some compounds that did not (C group, 27514, 31662) were affected by multiple exposures to light, with one (31662) surprisingly showing a very large increase (**Fig. 3b, Fig. S5**). Generally, the wavelengths at which peak absorbance occurred were most affected by multiple irradiation, but this was not always the case. It remains unclear if changes in absorption, whether increasing or decreasing, are linked to the identified behavioral phenotype groups; for example, whether an increase in absorption resulting from prior violet light exposure would allow compounds not in groups A or B to induce A- or B-type activity.

Finally, the majority of hit compounds and optovin, as well as the DMSO control, did not demonstrate notable maximum excitation or emission wavelengths (**Table 1**). Interestingly, outlier compounds were all found to be in either Group A or Group B. This suggests that there may be two different mechanisms behind the optovin-like response - one that involves photoreactivity, and one that occurs in the absence of it. For the Group C and Group D compounds, however, photoreactivity does not seem to be important.

### Phototoxicity is wavelength dependent

We noticed that following treatment, exposure to different wavelengths had a marked effect on toxicity for many of the compounds. This effect was observed most strongly following violet and blue wavelength exposure, which significantly increased toxicity in many cases. This suggests that these compounds are inert or mostly inert until activation by specific wavelengths. No compounds demonstrated significant increased toxicity following exposure to green or red wavelengths (compared to the no stimulus control), and baseline toxicity was observed in only two compounds (increased toxicity in the no stimulus control compared to the no stimulus DMSO control). Intriguingly, in one compound that displayed baseline toxicity (30580), exposure to red light actually increased survival time, suggesting a protective effect that warrants further investigation in future studies. These data suggest that the energy provided by violet or blue light is necessary for the conversion of these compounds to their active form, which then allows them to bind to receptors and induce an effect. Potentially, when these receptors are continually activated for long periods of time, their effect eventually becomes toxic. As many of the compounds that demonstrated increased toxicity were shown to be TRPA1b dependent, it is therefore possible that this receptor is TRPA1b, and experiments utilizing TRPA1bKO fish will help to confirm this and to determine the mechanism of toxicity.

### Optovin-like hits are TRPA1 dependent

In this study, TRPA1b dependence was observed in many of the A and B compounds, but not in any of the C and D compounds. To confirm TRPA1-dependence, we developed a line of TRPA1b KO zebrafish and assessed if their activity matched that of wild-type fish following treatment and stimuli exposure. Additionally, to determine if any of the induced behaviors were vision-dependent, we obtained zebrafish with a mutation in *atoh7*, which causes the complete loss of retinal ganglion cells and results in completely blind fish. Of the 14 A and B group compounds, 8 demonstrated a significant loss of phenotype in the TRPA1b KO fish, with several others approaching significance. However, of the Group A and B compounds tested in ATOH7 KO fish, loss of vision did not have an effect on this behavior. Conversely, TRPA1b dependence was not observed in any of the C and D compounds. However, of the compounds tested, loss of vision resulted in loss of the phenotypes induced by three compounds in this group (C: 15951, 23002; D: 27514). Taken together, these results indicate that while some compound induced behaviors, especially those most similar to optovin-induced behaviors, are non-visual and are mediated by TRPA1b photosensation, others are acting through different receptors and mechanisms. Of these, the C type compounds are of special interest; while they share with optovin the ability to induce light activated behaviors, they differ greatly in phenotype, wavelength response, vision dependence, and target.

### Future directions

While this study has revealed that a wide diversity of structures, photochemical properties, behaviors, and toxicities exist among light--dependent TRPA1 agonists, further research is needed to better understand both the mechanisms and the research or therapeutic potential of these compounds. Further structural analyses would help to elucidate the chemical moieties and energy changes responsible for light activation, allowing for the development of purpose-build analogues. Likewise, a better understanding of receptor-compound kinetics and the mechanisms of the hit compounds would help to answer questions about where and how the compound is binding, whether light activation occurs at the receptor-compound complex or on the compound alone, and whether any intermediates or downstream signaling pathways of TRPA1 play a role in the induced behaviors. To this end, patch clamp experiments could help to reveal patterns of channel opening and the location of binding, while mutations in potential TRPA1 binding sites could confirm those locations^43^. The presence of intermediates such as reactive oxygen species (ROS) could be detected by co-treating with DABCO, a ROS quencher, or by using a singlet oxygen indicator dye such as singlet oxygen green^42^, while bleaching or “pre-activation” of compounds would help to determine if exposure of only the compound to light is sufficient to induce the observed phenotypes. To ascertain the activity of these hit compounds on an organism-wide level, experiments on headless fish (as described^42^) could identify whether the compounds act in the brain or in the periphery, and if in the brain, pERK imaging could identify the specific location, while mass spectrometry analysis could reveal the neurotransmitters affected. Finally, it is crucial to determine whether these results carry over into other species. Mouse studies would reveal the effect of these compounds in a mammalian model, and comparisons to the effects of known TRPA1 agonists such as AITC could reveal key differences in their action. Furthermore, experiments on human TRPA1 transfected cells would confirm the compounds’ ability to induce an effect in humans, and thereby open the door for potential medical research or therapeutic use.

## METHODS

### Fish maintenance, breeding, and compound treatments

Eggs (up to 10,000 per day) were collected from group matings of wild-type “Singapore” zebrafish, and raised until 7dpf on a 14/10-hour light/dark cycle at 28°C in egg water (60 µg/ml “Instant Ocean” Sea Salts in distilled water, then buffered and brought to a pH of 7.0-7.4 using sodium bicarbonate^44^). Following sorting for healthy zebrafish and temporary anesthetization using egg water chilled to 4°C, eight larvae and 300µL of egg water per well were distributed into square 96-well plates (GE Healthcare Life Sciences) using a p1000 pipette as described^29^. Plates were allowed to return to room temperature over 1 hour, after which compounds diluted in DMSO were applied directly to the egg water. Larvae were incubated following treatment for an additional hour before behavioral analysis. All zebrafish procedures were approved by the UCSF’s Institutional Animal Care Use Committee (IACUC) in accordance with the Guide to Care and Use of Laboratory Animals (National Institutes of Health 1996) and conducted according to established protocols that complied with ethical regulations for animal testing and research.

### Compound preparation and chemical libraries

Chemical libraries were selected and purchased from the Chembridge DIVERSet Screening Libraries. All chemical library compounds were stored at -80C and dissolved in DMSO, then further diluted in DMSO and screened at a final concentration of 10µM in <1% DMSO. All controls were treated with an equal volume of DMSO. Hit compounds were obtained from commercial sources (Hit2Lead, TimTec, Molport, Vitas, Mcule, ChemDiv). Optovin was obtained from Tocris (4901), DMSO from Sigma Aldrich, AITC from Sigma Aldrich (36682-1G), and cinnamaldehyde from Sigma Aldrich. Following receipt, compounds were initially diluted to 30mM stock solutions in DMSO and stored at -20°C. Immediately prior to treatment, compounds were thawed at room temperature and further diluted with DMSO as required.

### Automated behavioral phenotyping assays

Phenotypes were induced and recorded using a custom-built instrument that includes a camera, lighting, and various light, audio, and physical stimuli, as described in Myers-Turnbull et al. 2020^45^ and McCarroll et al. 2019^29^. Briefly, plates were illuminated with a 760nm infrared light, and an AVT Pike digital camera (Allied Vision) fitted with an infrared-passing/visible light-blocking filter (LEE Filters LE8744 polyester #87) and mounted to a telecentric lens captured images at 25fps. Light stimuli were delivered using high-intensity LEDs and included red at 623nm (1537-1041-ND, DigiKey), green at 525nm (1537-1039-ND, DigiKey), blue at 460nm (1537-1037-ND, DigiKey), and violet at 400nm (LZ4-40UB00-00U7, Mouser). Acoustic stimuli were delivered using a computer to playback audio stimuli as MP3s using an APA150 150W powered amplifier (Dayton Audio) played through surface transducers adhered to the acrylic stage. Physical tapping stimuli were delivered using push-style solenoids (12V) to tap a custom built acrylic stage. Instrument control, including data acquisition and stimulus activation, was conducted using custom MATLAB and Python scripts. Processing of video data to the zebrafish motion index (MI) was calculated as follows: MI?=?sum(abs(frame_n_ – frame_n−1_)).

### Optovin and TRPA1b agonist activity analysis

Optovin phases were identified from data collected during a single, 10 minute long purple stimulus. Phases were defined as follows: Pre-stimulus (“pre”) begins at the start of the battery and ends at stimulus start. The average of this phase is also the optovin baseline. Phase 1 (“P1”) begins at stimulus start and ends following the occurrence of the lowest MI within the first 30s of response. Phase 2 (“P2”) begins following P1 and ends following the occurrence of the first MI value that is less than or equal to the optovin baseline. Phase 3 (“P3”) begins following P2 and continues through the end of the stimulus. Post-stimulus (“post”) begins following the end of the stimulus and continues through the end of the battery. Average MI values are averages throughout each phase for each individual well, which are then averaged across all wells. Overall optovin activity in response to different stimulus wavelengths is shown as AUC, calculated as the sum of all MI values during the sixth one minute stimulus for each well, then averaged over all replicate wells. Optovin activity in comparison to known TRPA1 agonists is displayed as fold change vs. DMSO for each phase. Phases for all conditions are defined by optovin activity (as above), and values calculated by averaging MI over each phase for each well, then averaging data across all wells. Fold change was calculated by dividing each of these values by the average DMSO value. Significance was calculated vs. optovin using Welch’s t-test.

### Compound screen hit identification and hit retest analysis

To identify first-level hits from screened library compounds, “regions of interest” (ROIs) containing violet wavelength stimulus were identified for each assay within the full battery. Average MI values from each ROI were then obtained for all compounds. Fold change vs. DMSO was then calculated by dividing values from each ROI for each compound by the averaged data from the equivalent DMSO control ROI from the same plate. A fold change value for each DMSO well was calculated in the same way, with ROIs from each DMSO on a plate divided by averaged data from the equivalent ROI for all other DMSO controls from the same plate. The maximum fold change value from each compound and DMSO was then selected and plotted as a histogram. Compounds that had a maximum fold change greater than +3σ from the mean were selected for further analysis. Following initial identification, hits were further filtered by several levels of analyses. Manual analysis of compound screen MI traces and video removed any anomalous traces, after which retesting at 10µM against the screening battery with three replicates identified compounds with reproducible phenotypes. Compounds that reproduced were then selected for dose-response testing at 8 doses between 0.78125µM and 100µM (x6 replicates) against the screening battery. ROIs were again selected, and average MI values calculated. Significance for ROIs from each compound was calculated vs. same-plate DMSO wells using Welch’s t-test. A dummy DMSO significance value was obtained by randomly splitting DMSO replicates from each plate into the “test” and “control” groups. The significance threshold was determined after applying the Bonferroni correction (0.05/3014 tests).

### Photochemical analyses

Hit compounds and optovin were diluted to 100µM in egg water. An equal volume of DMSO in egg water and plain egg water were used as controls. To measure excitation/emission wavelengths, a 3D Fluorescence Scan was performed on a Tecan Spark. Compounds were plated onto a Grenier 384 well plate (Sigma, M6811). Emissions were measured every 20nm from 375-735nm, and excitations were measured every 20nm from 350-690nm. To measure absorbance, an Absorbance Spectrum experiment was performed on a SpectraMax M5 Microplate Reader (Molecular Devices). Compounds were plated onto a 384 well UV Star plate (Grenier Bio-One, 781801). Absorbance wavelengths were measured every 20nm from 250-750nm at 25C. For the multiple irradiation experiments, compounds were plated, then placed into the custom phenotyping instrument with the violet light stimulus activated. Immediately after light exposure, absorbance was measured as described above. Compounds were exposed to the violet light for varying lengths of time between 5 and 1200s and were freshly plated for each exposure.

### Phototoxicity assay

Fish were incubated with hit compounds for 1 hour prior to stimulus exposure. After incubation, fish were placed into the behavioral phenotyping instrument and their activity tracked for 12 hours. During the first six, larvae were exposed to a light stimulus at one wavelength (blue (460nm), violet (395nm), red (660nm), green (523nm), or no stimulus), after which activity was tracked for an additional six hours. Time of death was determined by identifying the 5 minute window in which the standard deviation of activity (as measured by MI) was equal to or less than the overall standard deviation of activity in the 300μM eugenol-treated positive controls. Additionally, the standard deviation of the remainder of the MI trace (beginning after the 5 minute window) could not exceed the eugenol standard deviation; if it did, the next 5 minute window was selected and tested as above. Wells that did not meet these criteria within the 12 hour period were considered to have survived. Average length of survival was calculated by averaging time of death across all wells treated with the same compound and exposed to the same stimulus. Significance was calculated using Welch’s t-test. For the violet, blue, red, and green conditions, the same compound with no stimulus exposure was used as the control. For the no stimulus condition, DMSO with no stimulus exposure was used as the control. A p-value of 0 indicates no difference from the control.

### Structural and phenotypic clustering

Structural clustering was performed with custom Python scripts using the SciPy hierarchical clustering (fcluster, linkage) and distance (pdist, parameters: metric = ‘jaccard’) functions. Phenotypic t-SNE analysis was performed on full MI traces with custom Python scripts using the scikit-learn t-SNE function (parameters: perplexity=10, learning_rate=500, n_iter=3000, n_iter_without_progress=500). Pairwise comparisons of selected features were performed by first selecting features of interest and calculating the relevant values for each compound, including phases “pre”, P1, P2, and “post” (individually) vs. DMSO fold change, P1 and P2 (individually) vs. baseline fold change, and average MIs for each individual phase. Each of these values was then plotted in a pairwise fashion and analyzed manually to identify the features that generated the greatest separation. Heatmaps were plotted by averaging MI values by second and using the Seaborn clustermap function. Final phenotypic classifications were determined manually. Following classification, phenotypic groups were retroactively labelled on the t-SNE analysis and pairwise feature comparison to verify that manual classification was in agreement with quantitative clustering.

### TRPA1b KO and ATOH7 KO zebrafish line development

The TRPA1b KO zebrafish line was generated using CRISPR to target a sequence (5’ CTGCACTATGCTTCAGCTGG 3’) in exon 3 of the *trpa1b* gene. Injections were performed as described in Burger et al. 2016^46^. The F0 generation was allowed to mature into adults, then crossed with WT TUAB zebrafish. The resulting F1 generation was checked for the presence of a mutation using PCR and restriction digest with PvuII. DNA from fish with a mutation was then sequenced to confirm that the mutation introduced a stop codon, and F1s that had the same mutation were crossed together. From the F2 generation, TRPA1b -/- fish were selected and crossed, resulting in an F3 generation consisting entirely of TRPA1b -/- fish. To obtain larvae for experiments, F2 or F3 TRPA1b -/- fish were placed in breeding tanks with males and females separated the night before crossing. The next morning, the divider was pulled, and any resulting embryos were collected. Larvae were used for behavioral experiments at 7dpf. ATOH7 zebrafish larvae were generously provided by the Schoppik lab at NYU. Once the larvae were mature, fish were genotyped using PCR and restriction digest with StuI, and ATOH7 +/- fish were crossed together. ATOH7-/- larvae, which are visibly darker than ATOH+/+ or +/- fish, were then selected and raised. Because ATOH7-/- zebrafish are blind, it is necessary for them to be raised separately from homozygous or heterozygous fish to prevent out—competition for food, and it is critical that for the first two weeks, the fish are provided with a high density of live food. To this end, starting at 5dpf, ATOH7-/- fish were raised in a separate tank and provided with at least 150mL live paramecia per day. From 14dpf to 20dpf, fish were fed with both live paramecia and dry food (Gemma 75). Beginning at 21dpf, fish were fed with dry food only. Larvae for experiments were obtained by crossing ATOH7-/- fish and following the breeding and collection protocol above.

### TRPA1b KO and ATOH7 KO zebrafish analysis

To assess whether the novel hit compounds depend on the TRPA1b receptor to induce a phenotype, WT and TRPA1b KO zebrafish larvae (five fish per well^45^) were treated with the same compounds and the activity response compared. Following selection of regions of interest (P1, full 1 minute stimulus, first three 1 minute stimuli), the mean MI for each of these ROIs was calculated. Fold change vs. control was then determined by dividing by background-matched DMSO controls. Significance between WT and TRPA1b KO responses was determined using Welch’s t-test. To assess whether vision is required for the hit compounds to induce a phenotype, WT and ATOH7 KO fish were compared in a similar fashion. However, because control variables were too variable, mean MI values were used for comparison and to determine significance rather than fold change.

## AUTHOR CONTRIBUTIONS

Conceptualization, D.C. and D.K.; Methodology, D.C., M.N.M., and J.C.T.; Investigation, D.C., M.N.M, and T.W.; Software and Formal Analysis - D.C.; Writing - Original Draft, D.C. and D.K.; Writing - Review and Editing, D.C. and D.K.; Supervision, D.K.

## ACKNOWLEDGEMENTS

The authors would like to thank Douglas Myers-Turnbull, Chris Li, and Cole Helsell for their assistance with coding, data management, and hardware development, Amanda Carbajal for experimental and technical support, and Louie Ramos, Nolan Wong, Madison Burns, Vy Nguyen, and Alexis Marquez for animal husbandry. Thanks also to David Schoppik at the NYU Langone Medical Center for generously providing the ATOH7 -/- zebrafish and recommendations on their husbandry. Finally, thanks to Jason Gestwicki and Su Guo for providing feedback and advice on experimental design and direction. Funding was provided by R01AA022583 (DK), and Pharmaceutical Sciences and Pharmacogenomics Training Grant GM008284.

## GRAPHICAL ABSTRACT

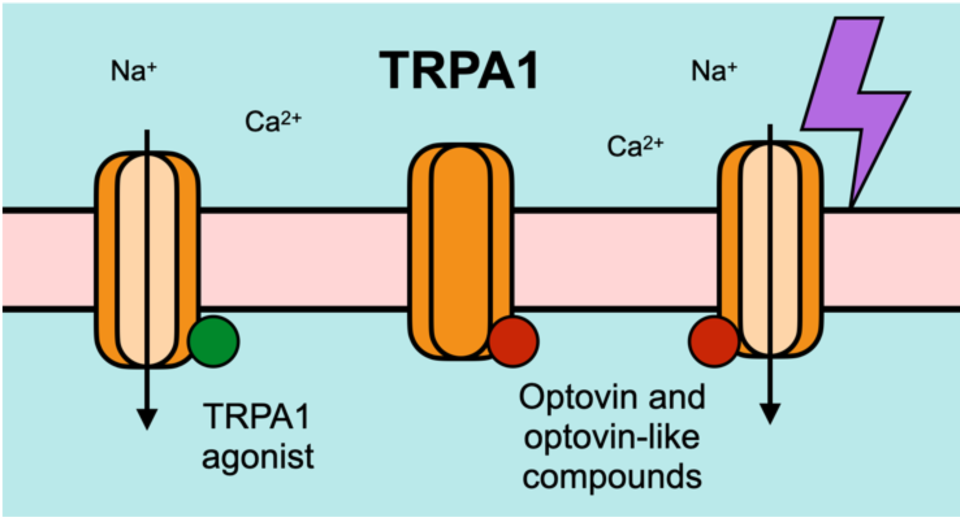

